# Fiber-based hydrogels for designing viscoelastic responses in particle-based biomaterials that support embedded 3D printing

**DOI:** 10.1101/2025.06.09.658654

**Authors:** M. Gregory Grewal, Lauren Porter, Georgia T. Helein, Christopher B. Highley

## Abstract

Viscoelastic biomaterials that exhibit biomimetic responses to applied stresses are important in studying physiology and designing biomaterial scaffolds. Particle-based hydrogels offer potential for engineering viscoelasticity through design of both the component microparticles and their processing into bulk particle-based materials. When particles are not crosslinked to one another, particle movements in response to strain can potentially relieve applied stresses and facilitate the material’s use in dynamic processes like bioprinting. In particle-based hydrogels based on spherical hydrogel microparticles (HMPs), particle movement is restricted by contacts with immediately adjacent HMPs. In comparison, fiber-based hydrogel systems leverage high-aspect ratio microfiber components with long-range interactions. Here, microfibers with aspect ratios of ∼15:1 length:diameter are used to form particle-based hydrogels to compare how interparticle interactions at increased length scales alter properties compared to particle-based hydrogels based on spherical HMPs. Like particle-based hydrogels formed from spherical HMPs, those formed from fiber HMPs exhibit viscoelasticity, with shear-thinning and self-healing behaviors. But, fiber-based materials allow enhanced control over bulk stress relaxation times (T_1/2_ ∼ 1-100+ s) across a range of applied strains (σ ∼ 2.5-50%) in a packing density-dependent fashion. Fiber-based systems relaxed stresses continually and to a greater degree at low strains in comparison to HMP systems. Dynamic interfiber interactions in fiber-based hydrogels also supported embedded printing where perfusable channels can be printed into fiber-based hydrogels stabilized by physical interfiber interactions. Taken together, fiber-based hydrogels offer opportunities for designing complexity into biomaterial scaffolds, including allowing control over viscoelastic properties through hydrogel design and control over heterogeneous 3D structures through embedded printing.

## I. Introduction

Biomaterials for tissue engineering and *in vitro* models of physiology and disease aim to recapitulate the complex viscoelastic mechanics of native tissue extracellular matrices (ECMs)^1–3^ that are necessary for faithful biomaterial models of tissue systems^4–7^. Dynamic mechanical ECM responses to cellular activity are understood to drive fundamental cell behaviors^8,9^. Permissive, yielding material environments are influential in proliferation, stem cell maintenance of differentiation, migration, and organization of multicellular structures^6,7,10–13^. Accordingly, many hydrogel biomaterials aim to recapitulate viscoelastic features^14,15^ of the ECM including complex time-dependent responses of the ECM to applied stresses and strains^16–20^.

Hydrogels can be designed to match many of the biophysical and biochemical attributes of natural tissue, with 2D and 3D systems advancing knowledge of cell-material interactions and the influence of environmental properties on cell fates^14–18,21,22^. Cell-material interactions in 3D environments are often more complex than in 2D systems^22^, and research designing 3D hydrogel systems looks to offer alternative platforms to natural materials like collagen, fibrin, and decellularized ECM-derived materials that are often used in studying cellular responses within viscoelastic and stress relaxing 3D environments^2,3,13,23,24^. Natural materials typically offer limited opportunity to controllably define biochemical and biophysical properties. Additionally, these materials may vary from batch-to-batch and may present low moduli with limited potential for designing biophysical features, which can limit their utility in engineered systems^9,25^.

These challenges have inspired the development of hydrogel systems that enable the design of complex mechanical properties. Hydrogel permissiveness to applied forces through molecular network yielding can be achieved through the design of degradable polymeric networks, but has also been enabled through engineering molecular structures to achieve dynamic, stress yielding materials^14,16,26^. Examples of material design to engineer viscoelasticity and stress relaxation into hydrogels include dynamic guest-host chemistries grafted onto various polymer backbones^27,28^, physical (ionic) chelation of polymers like alginate^29,30^, reversible covalent chemistries^12,31^, and using peptide interactions designed to modulate dynamic behaviors in a hydrogel’s nanofibrous network^32,33^.

Permissiveness and yielding can also be designed in particle-based hydrogels, which have intrinsic dynamic yielding and permissive properties, and there has been considerable work developing these 3D biomaterials^34–36^. These hydrogel systems are comprised of discrete hydrogel microparticles (HMPs) that are packed together. While interparticle crosslinking, or annealing, can be used to stabilize particle-based hydrogels against shear^37^, HMP-based materials can maintain a solid-like state by contact forces between discrete particles alone^34–36,38,39^, with dynamic rearrangements of HMPs possible in unannealed systems when sufficient force is applied^40–42^. In comparison to continuous hydrogels, where yielding occurs through rearrangements in the molecular network, yielding HMP systems occurs at longer length scales dictated by particle sizes. Additionally, the micro-to-mesoscale space among HMPs inherently enables cells to migrate through the scaffold^43–45^.

While particle-based hydrogels are typically formed from spherical HMPs, hydrogel microfibers in packed aggregates form hydrogel bulks that might be stabilized by designed interfiber chemical bonds or physical interfiber interactions, such as entanglements and jamming^46–48^. Within unannealed particle-based hydrogels, the dynamic rearrangements of HMPs that can occur in response applied forces are different in the case of microfiber-based HMPs, compared to spherical HMPs, because of high-aspect ratio particle morphologies. Fiber-based particles in closely packed systems can participate in longer range interactions with more particles than spherical HMPs and also have the potential to entangle and bend. These features can both lend stability to the system^46–48^ but also allow for microscale reorganization of fibers in response to applied stresses, including cell-scale forces that are sufficient to reorganize synthetic fibrous matrices^49,50^.

Recently, work by our group and others has considered how the properties of 3D microfiber-based materials might be useful in biomedical applications, ranging from forming 3D cellular microenvironments^50,51^, to bioprinting^47,51,52^, to providing scaffolds for tissue regeneration^47,48,53,54^. Among the unique physical properties that motivate many of these microfiber-based HMP applications are stress relaxation on time scales of 10s of seconds^51^, as seen in biological tissues^13^. Here, we sought to understand how dynamic behaviors in microfiber-HMP particle hydrogels compared to those in spherical-HMP based hydrogels, how changing the hydrogel design might affect bulk-stress relaxation responses, and whether microfiber-HMPs might serve as a support material for embedded 3D bioprinting. We report on a fiber-based particle hydrogel system, engineered from polyethylene glycol (PEG)-based microfibers, whose biochemical and biophysical properties have been shown to be highly designable^55,56^. We compare fiber-based particle hydrogels to those based on spherical particles and show that the former exhibits tunable viscoelasticity and stress relaxation responses, and that these responses are distinct from spherical PEG-microgel particles. Additionally, the dynamic behaviors and long range interparticle interactions within these PEG-fiber systems support embedded printing, opening possibilities diverse potential applications in engineering biomaterials with complex properties.

## II. Materials and Methods

### PEG microfiber production

Microfibers were formed by electrospinning a polyethylene glycol (PEG)-hydrogel solution. This electrospinning solution consisted of 10% w/v PEG norbornene (PEGNB, 20 kDa, 8 arm, JenKem Technology), 7% w/v PEG thiol (PEGSH, 10 kDa, 4 arm, JenKem Technology), 5% polyethylene oxide (PEO, 400 kDa, Sigma Aldrich), and 0.05% w/v 2-hydroxy-4’-(2-hydroxyethoxy)-2-methylpropiophenone (HHMP, Sigma Aldrich) in deionized (DI) water, which was dissolved overnight prior to electrospinning. The relative concentrations of PEGSH to PEGNB were used to create a stoichiometric mismatch between norbornene and thiol functionalities, in order to leave unreacted norbornenes for coupling thiolated peptides. 0.5 mM of a fluorescein amidite (FAM)-conjugated peptide containing a cysteine (GCDDD-FAM) for conjugation to free norbornenes was included in the electrospinning precursor solution when fluorescent-labeling was desired to enable imaging of microfibers. To reduce interfiber bonding between fibers during post-electrospinning crosslinking, a second solution containing 5% w/v PEO (900 kDa, Sigma Aldrich) dissolved in DI water overnight was prepared for co-spinning sacrificial fibers.

The PEG and sacrificial fiber solutions were both extruded through 16-gauge needles positioned 18 cm away from opposite sides of a rotating mandrel collector at rates of 0.4 ml/hr and 0.5 ml/hr, respectively. A voltage of 11-12 kV was applied to the needle attached to the PEG solution, and 5.5-6.5 kV was applied to the needle attached the sacrificial fiber solution. The mandrel had a -4 kV applied voltage and rotated at 1000 RPM to align fibers and minimize interfiber welding. Fiber batches were collected for 1 hr, crosslinked while dry under nitrogen for 15 min at 5 mW/cm^2^ (VWR UV Crosslinker) to stabilize the PEG fibers, then immersed in PBS to hydrate PEG fibers while simultaneously dissolving sacrificial fibers. Fiber batches were hydrated overnight to ensure full swelling of the hydrogel network and removal of undesired components from the electrospinning process (PEO and unreacted photoinitiator).

Hydrated fiber mats were then suspended in PBS at ∼10% v/v, segmented via homogenization (IKA T25) at 10k RPM for 2 min, and filtered through a 40 *μ*m mesh to remove welded fiber aggregates. Fiber suspensions were centrifuged and resuspended thrice to remove all undesired components, before resuspending fibers 10% v/v (10% v/v indicates 1 part centrifuged fiber segments in 9 parts PBS) and stored at 4 °C until use. Fiber length and diameter were characterized using ridge detection in ImageJ based on thresholded confocal images of dilute fiber solutions (BioTek Cytation C10, Agilent Technologies).

### Peptide Synthesis

The fluorophore-conjugated peptide, GCDDD-FAM, was used to visualize microfibers and microgels in this study. The peptide was synthesized with a cysteine residue to permit thiol-ene conjugation to residual norbornenes during the fiber- and particle-making processes (Liberty Blue automated, microwave-assisted solid phase peptide synthesizer, CEM). The peptide was built from C-terminus to N-terminus on Rink amide resin using Fmoc-protected amino acids (resin and amino acids were sourced from Advanced Chemtech), with 5(6)-carboxyfluorescein (5(6)-flourescein amidite or FAM, Sigma Aldrich) added last to the N-terminus. The resultant peptide was cleaved off the resin using a cocktail of trifluoroacetic acid, triisopropylsilane, 2.2’-(ethylenedioxy) diethanethiol (all were sourced from Sigma Aldrich), and DI water at a 92.5/2.5/2.5/2.5 mixing ratio, respectively. The freed peptide was then isolated via precipitation in cold diethyl ether (Sigma Aldrich), dried under vacuum, resuspended in DI water, and lyophilized to yield the final product. Peptide synthesis was confirmed using MALDI-TOF spectrometry (Figure S1).

### Aqueous two-phase PEG hydrogel microparticle synthesis

PEG hydrogel microparticles were generated via aqueous two-phase suspension. A solution of dextran from *Leuconostoc* spp. (70 kDa, dextran(70), Sigma Aldrich) was mixed with a given PEG hydrogel precursor solution, as described below, at a 4:1 ratio of continuous (dextran) to dispersed (PEG) phases. For volume-matched spherical microparticles (spheres (V)), 800 *μ*L of 40% w/v dextran, and 0.05% w/v HHMP in DI water was mixed with 200 *μ*L of a solution comprised of 6% w/v PEGNB, 4.2% w/v PEGSH, and 0.05% w/v HHMP in DI water. When preparing particles for fluorescence imaging, 0.5 mM of GCDDD-FAM peptide was included in both phases. This mixture was vortexed at maximum speed for 1 min prior to crosslinking under UV light at 20 mW/cm^2^ (Omnicure) for 5 min. For volume-matched spherical microparticles (spheres (D)), the continuous phase was modified to contain 25% w/v dextran and the stir rate was reduced to 800 RPM, but all other parameters were conserved compared to spheres (V). Following crosslinking, the resultant particles were suspended in 15x volume of PBS to thermodynamically shift towards a single phase solution and centrifuged twice to remove dextran and other unreacted materials. Spheres (V) and spheres (D) were filtered through 20 *μ*m and 40 *μ*m meshes, respectively, and centrifuged once more to yield the final particles used within this study. Particles were also stored at 10% v/v suspension in PBS at 4 °C until further use. Similar to fibers, particle size was characterized using ImageJ based on thresholded confocal images of dilute particle solutions (BioTek Cytation C10, Agilent Technologies).

### Forming particle-based hydrogels

Two classes of particle-based hydrogels were designed for this study: those comprised of PEG microfibers and those comprised of spherical PEG microparticles. Fibrous particle hydrogels were assembled via centrifugation at 5k, 10k, and 15k RCF for 5 min to yield low, medium, and high packing densities, respectively. Spherical particle hydrogels were formed from either spheres (V) or spheres (D). These were packed at the medium packing density to enable comparisons with the fibrous assemblies packed at medium density. Following centrifugation of a given particle-based hydrogel, the supernatant was carefully aspirated to avoid disrupting the particle-based hydrogel and the pellet was manipulated using a spatula thereafter.

### Particle-based hydrogel characterization

To characterize void space in particle-based hydrogel assemblies, fibers and spheres were resuspended in 2 mg/mL FITC-dextran (2 MDa) prior to centrifuging to form particle-based hydrogels, then centrifuged to yield the desired packing density. Through microscopy, fluorescent signal would be seen within the void space of the particle-based hydrogel sample. After centrifugation, particle-based hydrogels were then transferred to a 96 well plate and Z-stacks of each sample were acquired at random ROIs on a Leica Stellaris 8 confocal microscope. Images were thresholded on ImageJ and void space was quantified using the built-in Analyze Particles functionality. Void space was determined as the average pixel intensity of the fluorescent regions with respect to the total pixel volume of the micrograph for each group.

Mechanical properties of particle-based hydrogel assemblies were assessed via oscillatory shear rheology (DHR-3, TA Instruments) using a 20 mm parallel plate geometry, a 500 *μ*m gap distance, and a 25 °C testing temperature. Time sweeps (0.5% strain, 1 Hz) were utilized to assess viscoelasticity of particle-based hydrogels. Cyclical application of low and high strains (low: 0.5%, 1 Hz; high: 250%, 1 Hz) were used to observe shear recovery. Strain sweeps (0.01%-500% strain) were used to elucidate strain yielding and critical strain values. Finally, a single application of a defined strain (ranging from 2.5%-50%) was utilized to investigate stress relaxation characteristics of particle-based hydrogel assemblies.

### 3D printing into fiber particle-based hydrogels

A removable biomaterial ink was formed from packed gelatin microparticles containing either a black dye (India ink) or 2 MDa FITC-dextran for visualization via either widefield or fluorescent microscopy. Gelatin microparticles were formed by heating a 15 w/v% solution of gelatin with India ink in PBS to 80 °C and a separate volume of mineral oil to 40 °C. Gelatin solution was then added to the mineral oil at a 1:10 ratio (aqueous:oil) and homogenized at 9,000 rpm while the emulsion cooled to room temperature. The particles-in-oil suspension was then cooled to 4 °C and centrifuge to pellet the gelatin microparticles. The oil supernatant was aspirated, then washed twice with isopropanol on a 0.22 µm PVDF membrane in a Buchner funnel via vacuum aspiration. Dried particles were collected, washed in 70% ethanol overnight. Prior to use, gelatin microparticles were rehydrated in PBS and centrifuged to form a gelatin microparticle-based hydrogel, which was loaded into a syringe for printing. Gelatin microparticle sizes were quantified via ImageJ from widefield images.

To prepare fiber particle-based hydrogels as support materials for embedded printing, fiber-based hydrogels prepared at the highest packing densities were transferred into wells within a polydimethylsiloxane (PDMS) device. Printing was performed using a FELIX Bioprinter that was equipped with positive displacement printheads. Print paths and extrusion were defined via directly written G-code commands. G-code commands were executed to deposit filaments of gelatin microparticle ink into the fiber particle-based hydrogel support. The ends of the filaments aligned with channels that could introduce fluid into the center well of the PDMS device via hydrostatic pressure. After printing, the constructs were warmed to 37 °C to melt the gelatin, allowing media to enter the space that was occupied by the filament. Fluorescent polystyrene beads suspended in the media were used to visualize channels into which the media flowed.

## III. Results and Discussion

### 3.1. Preparing particle-based hydrogels

Both microfiber- and spherical microparticle-based hydrogels were generated using polyethylene glycol (PEG) in order to develop hydrogel systems that might be engineered to specific biomedical applications through design of chemical functionalization, polymer concentration, and crosslinking. Here, we formed microfibers by electrospinning solutions of PEG-norbornene (PEGNB) and PEG-thiol (PEGSH), a formulation that can be stoichiometrically designed to leave unreacted norbornene moieties^55,56^. This allowed additional thiol-ene conjugation of a thiol-containing fluorescent peptide in work here, but could support conjugation of bioactive peptides in applications with cells^55–58^. High-aspect ratio microfibers were formed by electrospinning the PEG hydrogel precursor solution (Figure 1A-B) and segmenting the fibers collected (Figure 1A-B). The process used here homogenized fiber suspenstions at 10k RPM for 2 min, then filtered suspensions to remove aggregates. The hydrated fibers formed through this process were approximately 37.3 *μ*m in length (Figure 1C) and 2.41 *μ*m in diameter (Figure 1F). This gave an aspect ratio (L/D) of ∼15, which was large compared to spherical particles with aspect ratios of 1.

**Figure 1.**
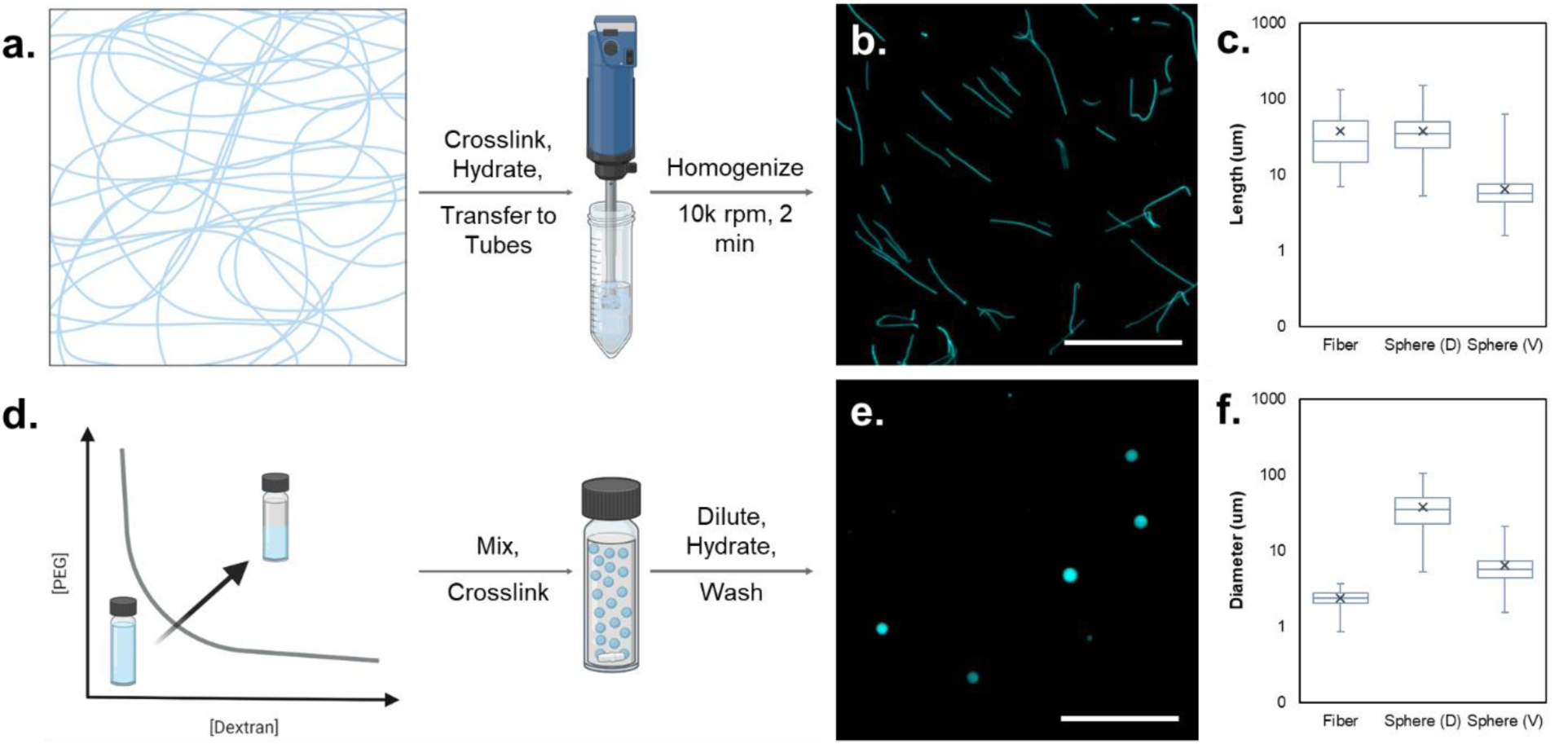
Preparation of particles for forming bulk hydrogels. (a) Electrospun PEG fibers were crosslinked, hydrated, and homogenized to segment fibers in a fast, scalable fashion. (b) Fluorescent micrograph of segmented PEG fibers. (c) Quantification of fiber and sphere length illustrating matching of fiber and sphere dimension in spheres (D) group. (d): Schematic of a binodal curve for PEG and dextran, where two phases occur in the regime above the curve. The ATPS system was then mixed to form dispersed PEG spheres within the continuous dextran phase, the hydrogel microparticles were crosslinked, diluted to form a single phase, and finally washed. (f) Quantification of fiber and sphere diameter illustrates the disparity in the dimensions between fibers and spheres. The aspect ratio of spheres was assumed to be ∼1, so length and diameter was assumed to be equal for these groups. Scalebars in b and e = 200 *μ*m; n > 300 for all groups.

To compare particle-based hydrogel properties between materials composed of fiber-based and spherical particles, spherical hydrogel microparticles were also prepared. Here, we used an aqueous two-phase separation technique^59–62^ (ATPS, Figure 1D-E) to yield batches of spherical particles with either matched lengths (particle diameter ≈ fiber length) or matched volume, where spherical volumes were assumed for particles and cylindrical volumes for fibers. These groups are referred to as Spheres (D) (for matched diameter) and Spheres (V) (for matched volume), respectively. ATPS leverages PEG-rich and dextran-rich aqueous solutions that are thermodynamically immiscible when mixed at high enough concentrations^59^. A systematic approach to determine experimental concentrations of PEGNB/PEGSH and dextran solutions was leveraged to form PEGNB hydrogel microparticles (Figures S3-S5 provide a summary of different processing variables). Spheres (D) had an average diameter 37.2 *μ*m (Figure 1C and 1F, same value) which matches the average length of the electrospun fibers, and Spheres (V) had an average diameter of 6.4 *μ*m (Figure 1C and 1F, same value) which approximately matches the volume of the electrospun fibers.

### 3.2. Influence of particle size and shape on particle-based hydrogel microporosity and yielding

Particle-based hydrogels were formed via centrifugation-mediating packing at different speeds to yield low, medium, and high packing densities, in groups referred to as Low-Fiber, Med-Fiber, and High-Fiber, respectively. Spheres (D) and Spheres (V) were packed at the medium density only (Med-Sphere (D) and Med-Sphere (V)) for comparison to fibers. To investigate differences in porosity of the particle-based hydrogels, particles were packed with high-molecular weight FITC-dextran and confocal microscopy was used to visualize the fluorescent signal within pores (Figure 2A). Consistent with previous findings in other particle-based hydrogel systems^35,63^, increasing the packing density of fibers resulted in decreased void space (from 21% for Low-Fiber to 15% for High-Fiber). Both Med-Sphere (V) and Med-Sphere (D) possessed larger quantities of void space in the particle-based hydrogel (22% and 27%, respectively) when compared to Med-Fiber. Given the controlled packing force, this is attributed to small diameters and bending of the fibers allowing fibers to deform to fill void spaces during centrifugation. In comparison, spheres largely maintain their shape in packing, with limited deformation possible as soft materials.

**Figure 2.**
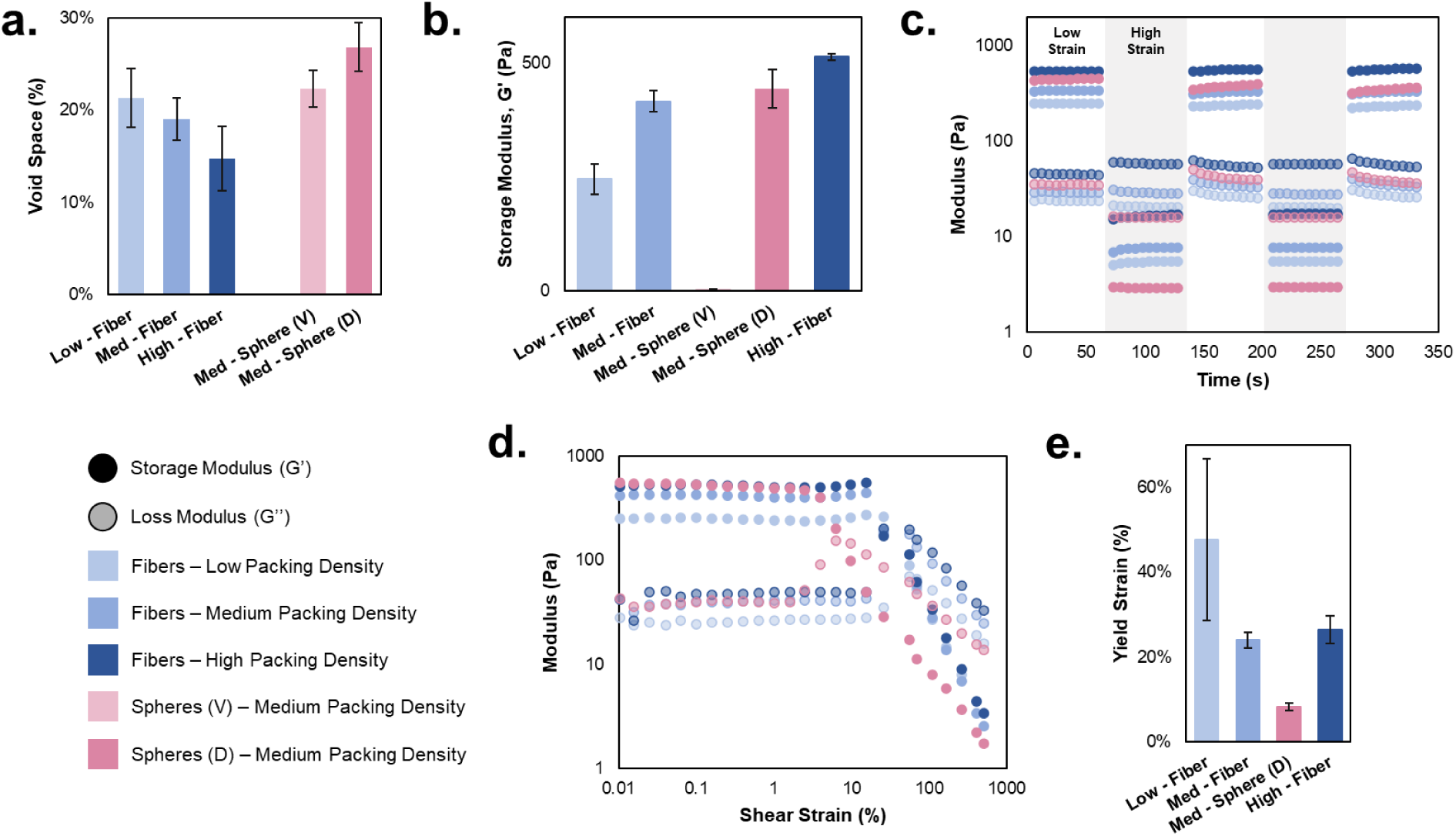
Particle size and shape influences overall particle-based hydrogel properties. (a) Particle-based hydrogels following centrifuge-mediated packing exhibit void spaces in both a packing density- and particle shape-dependent manner. (b) Storage moduli indicate that all groups exhibit solid-like behaviors at low strains, except for Med-Sphere (V) where there was no appreciable storage modulus, likely due to insufficient contact forces between particles. Generally, increased packing density yielded greater elastic contribution in particle-based hydrogels, with Med-Fiber and Med-Sphere (D) exhibiting similar stiffnesses due to similar characteristic dimensions defining the system.

In rheological measurements of mechanical properties, all fiber particle-based hydrogels behaved like elastic solids at low strain, with storage moduli ranging from ∼245 to ∼510 Pa (Figure 2B). Both Med-Fiber and Med-Sphere (D) exhibited storage moduli values of ∼450 Pa. Interestingly, the storage modulus measured for the Med-Sphere (V) was very low (< 5 Pa), which we attributed to their small, near-colloidal size. At colloidal scales, particles remain dispersed in suspension, and results here suggested that small size of the Spheres (V) group was likely limiting their packing. Consequently, only Spheres(D) were used in further analysis comparing bulk viscoelasticities of the particle-based systems.

Particle-based hydrogels lacking interparticle crosslinking can flow when sufficient stress is applied, allowing their use in injection^37^, extrusion^64^, or embedded printing^65^ as support matrices. Towards comparing these behaviors in the Med-Sphere(D), Low-Fiber, Med-Fiber, and High-Fiber groups, we exposed samples to cycles of low (0.5%) and high (250%) oscillatory strain. All systems demonstrated rapid transitions from solid-like to liquid-like behaviors and back upon the application and removal of high strain (Figure 2C). In using strain sweeps to compare where this transition occurred in each group (Figure 2D-E), we observed yield strains (% strain where G’’> G’) to be both packing density- and particle shape-dependent. More specifically, the yield strain (∼48%) for the Low-Fiber group trended higher than the yield strains for the Med-Fiber and High-Fiber groups (24% and 26%, respectively). We attribute this phenomenon to the higher interstitial fluid content in the Low-Fiber group effectively providing space for the fibers to move without altering interactions or entanglement. The spherical particle-based system (Med-Sphere (D)), exhibited a yield strain (∼8%) that was 50% lower than the lowest strain in the fiber groups. We expect that this difference results from spherical microparticles’ limited interactions within the particle-based system. Unlike microfibers, spherical microgels interact through near-neighbor interactions with no potential respond to strain through by extending out of entangled or bent conformations. Increasing strain is thus expect to force spherical particles to shift within the system, likely in brittle failure behaviors^66^. The comparatively enhanced ability of fiber-based hydrogels to support displacement while still exhibiting bulk-elastic behavior was expected to be a valuable addition to developing 3D biomaterials with permissive properties.

### 3.3. Stress relaxation in particle-based hydrogels

We next sought to compare how fiber- vs spherical-particles systems relaxed stress in response to applied and sustained strains. We expected that fibers’ interactions with an increased number of other fibers, compared to spherical particles’ interactions with next neighbors, and fibers’ potential for entanglements would allow them to sustain force over longer time scales in response to applied strain that could cause bulk flow. In other words, fiber-particles were expected to “slide” to reorganize microscale interfiber architectures through extension and relieving entanglement compared to spheres “shifting” en masse in response to an applied strain (schematic in Figure 3A) would result in strain relaxation over extended time in fiber-based systems, resulting in ECM-mimetic behaviors.

**Figure 3.**
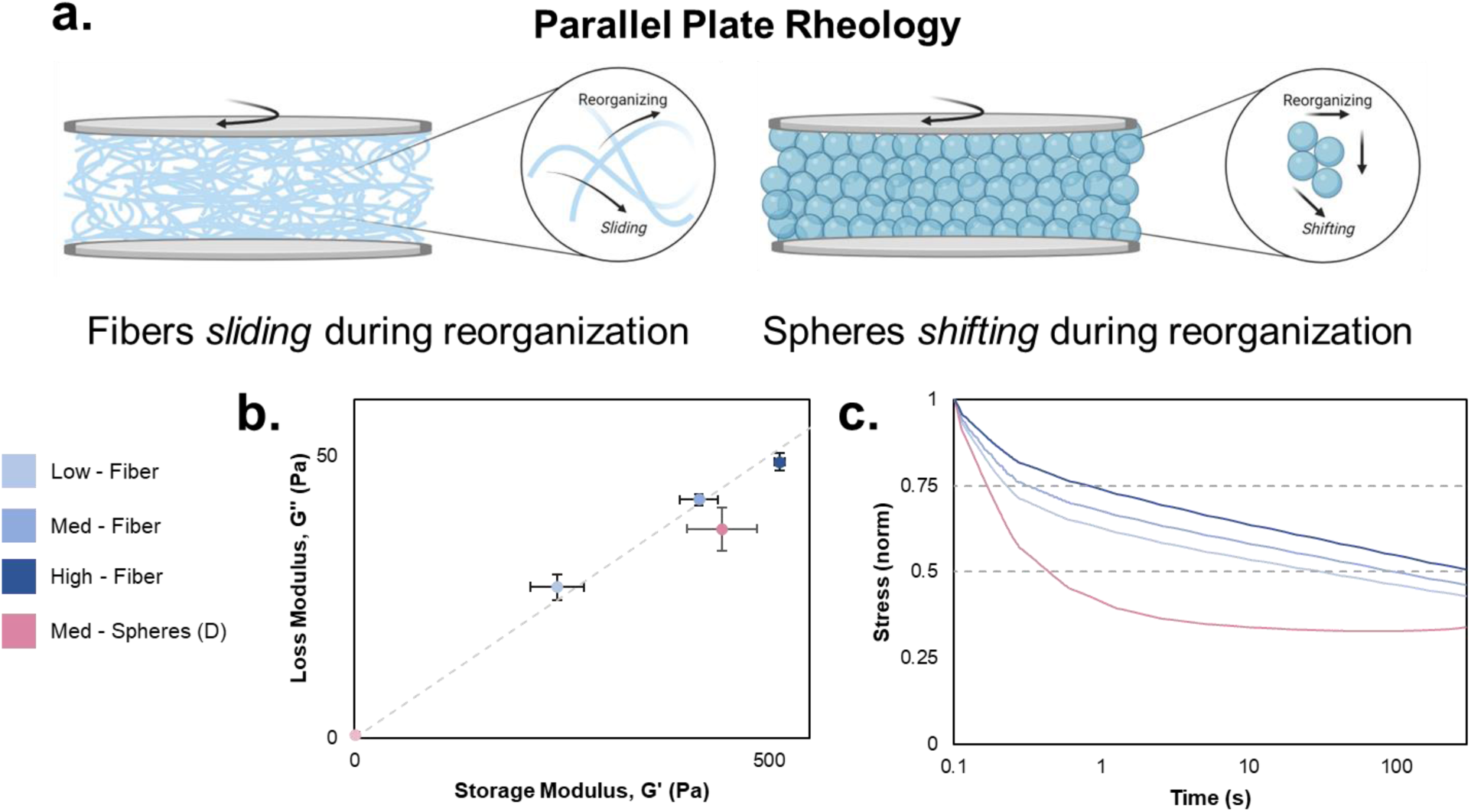
Viscoelasticity and time-dependent stress relaxation of granular hydrogels. (a) schematic of the parallel plate oscillatory shear rheology testing platform. The increased length scale of interactions between fibers enables sliding and reorganization in response to applied strains whereas particles will begin to shift in response to applied strains. (b) Plotting loss vs. storage modulus illustrates both viscous and elastic contributions to the mechanics of granular hydrogels. Fiber-based granular hydrogels exhibit storage moduli that are ∼10x loss moduli (illustrated by gray dashed trendline), which is consistent with many natural tissue types. Conversely, Med-Spheres (D) have a lesser viscous contribution and thus deviate from this 10x trend. Med-Spheres (V) illustrate negligible storage and loss moduli. (c) Time-dependent stress relaxation profiles at 15% applied strain of granular hydrogels. Fiber-based granular hydrogels are able to dissipate stress over time as fibers slide and reorganize in response to the applied strain as illustrated by the slow decrease in normalized stress (T_1/2_ on the order of 10-100+ s). Spheres (D) are unable to reorganize effectively and seemingly shift and fracture before reorganizing into a granular hydrogel as illustrated by the sharp drop in normalized stress (T_1/2_ < 1 s).

In considering biomimicry in fiber- and spherical-particle based hydrogels’ viscoelastic properties, both showed storage:loss moduli ratios of ∼10:1 (Figure 3B), which is characteristic of most tissues. In fact, the relative loss moduli contribution of the spherical-particle based hydrogels is comparatively smaller – giving a storage:loss ratio >10:1. This indicates that, in the fiber systems, enhanced microscale reorganization dissipates stress in oscillatory measurements. We next considered relaxation time, T_1/2_ – the time it takes for a tissue or material to relax to 50% of the peak stress under constant strain, typically 1-1000 s in biological tissues. Upon applying a constant applied shear strain of 15%, we found that all packing densities of fiber-based hydrogels were able to dissipate stress with T_1/2_ >10 s (Figure 3C). Additionally, T_1/2_ demonstrates a positive correlation with packing density, where the relaxation time is longest for the High-Fiber group. In comparison, Med-Spheres (D) exhibited a sharp drop off in normalized stress with a T_1/2_ <1 s. The relaxation times and profiles in the fiber systems more closely mirrored natural tissue, with the rapid drop in the spherical particle-based system suggesting the shifting of material en masse.

### 3.4. Strain-dependence of particle-based hydrogel stress relaxation

To further understand stress relaxation responses in the particle-based materials, we next looked at how material responses changed as a function of strain applied and, in the fiber-based hydrogels, packing density. We hypothesized that, if a strain were rapidly applied that exceeded yield strains, a spherical particle-based hydrogel would rapidly shift, while the relaxation response in a fiber-based system would be attenuated and prolonged as fibers reorganized. We applied a range of strains, 2.5 – 50%, to all systems (Figure 4A) to include strains both above and below measured yield strains (Figure 2E). For spherical particle-based hydrogels (Figure 4Ai), stress relaxation profiles evidenced two distinct dynamic regimes, as seen in profiles that differed based on whether applied strain exceeded yield strain (∼8% for the Med-Spheres (D), Figure 2E). In contrast, fiber-based materials exhibited stress relaxation profiles (Figure 4Aii-iv) whose shape indicate relaxation behaviors independent of yield strains (∼48%, 24% and 26% for Low-, Med-, and High-Fiber, respectively, Figure 2E). First, in the spherical particle group, for applied strains below the yield strain (<8%, Med-Spheres (D)), relaxation occurs gradually, with stress not reaching 50% of the applied stress after 300s. At applied strains of 10% and above, Med-Spheres (D) exhibit a sharp drop in stress held, followed by a relatively flat plateau, suggesting a shift of the particles that immediately relieves stress before stabilizing with some stress maintained, suggesting spherical particles are still strained, but below yielding.^41^

**Figure 4.**
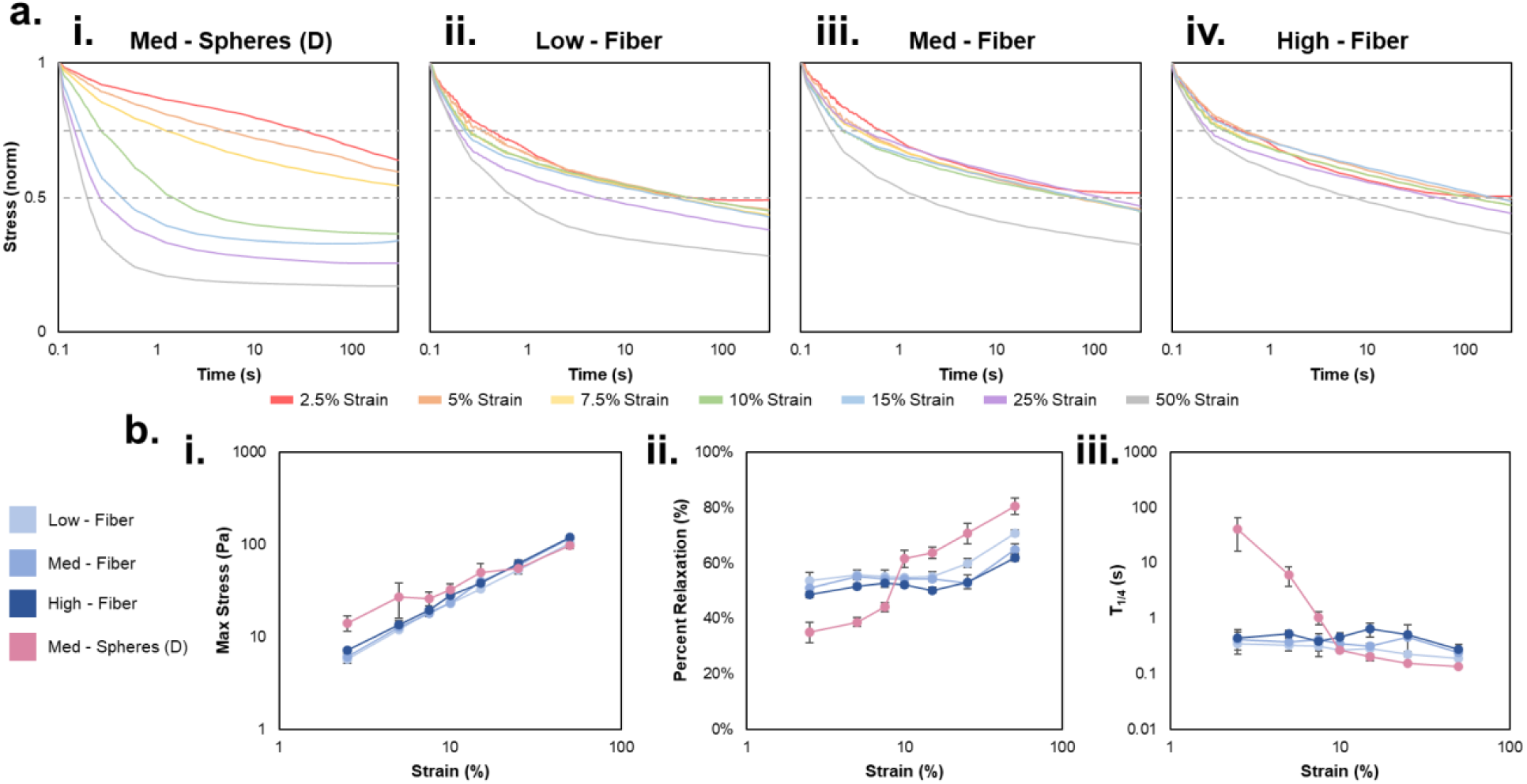
Strain-dependence of particle-based hydrogel stress relaxation. (a) Particle-based hydrogels formed from spherical particles, (i) Med-Spheres (D), exhibit yield strain-dependent relaxation profiles compared to (ii-iv) all packing densities of fiber-based hydrogels. Med-Spheres (D) reorganize and dissipate stress gradually below their yield strain (∼8%), whereas fiber-based granular hydrogels exhibit similar relaxation curves, across strain applied, including above yield strains. (b) This shift in behavior at yield strain that is seen in spherical particles but not in fiber particles, can be further quantitatively observed through (i) maximum stress for applied strain, with Med-Spheres (D) generally exhibiting an elevated stress at sub-yield strains; (ii) percent relaxation after 300 s, where Med-Spheres (D) do not relax to the same level as fiber-based groups at strains below their yield strain, but step to higher stress relaxation nabove yield strain; and (iii) Finally, consistent with the previous results of total relaxation, T_1/4_ is considerably longer for Med-Spheres (D) when the applied strain is below the yield strain, with a sharp decrease as the applied strain is increased beyond this threshold. Conversely, fiber-based granular hydrogels exhibit a marginal decrease in relaxation time as applied strain is increased.

In comparison, all packing densities of fiber-based hydrogels exhibit similar profiles that indicate applied strain is initial relieved through a rapid reorganization, followed by a gradual and continuing relaxation phase (Figure 4Aii-iv). As strains increase, stresses driving relaxation increase (Figure 4Bi), but relaxation only gets faster at the highest strains for any fiber group (Figure 4Bii). More porous fiber-based hydrogel also relax strain more quickly, indicating porosity, and decrease fiber interactions per volume, supports fiber reorganization. In considering T_1/4_, the time required to relax one-fourth of the stress resulting from an applied strain (T_1/4_ can be compared across all groups, whereas T_1/2_ was not reached in all groups under the tests here), these trends are also evident (Figure 4Biii). Fiber-based hydrogels show a relatively constant T_1/4_, that is interestingly similar across packing densities, and that decreases modestly with increasing strain. In the case of spherical particles, T_1/4_ decreases in the sub-yield strain regime, and is constant above the yield strain, in line with previously described behaviors.

Taken together, these results show that the fiber-based hydrogels described characteristically relax applied stresses on short time scales and in a strain-independent manner, compared to spherical particles. Additionally, the data show that fiber-based materials respond to high strains, and the corresponding increases in stress (Figure 4Bi) without large fracture-like shifts within their bulk, but rather through steady rearrangement of microscale fiber organization. Within the ECM, cells are known to exert protrusion and traction forces during migration on the order of 10^-1^-10^1^ kPa, coupled with 10-50% strains when they interact with their environment^41^. Fiber-based hydrogels therefore provide providing an opportunity to establish biomimetic 3D environments, given their combined low-strain stress relaxation and continued yielding responses at these strains.

### 3.5. Embedded printing within fiber-based hydrogels

Fiber-based hydrogels’ ability to flow and restabilize (Figure 2C) indicated potential to serve as a novel support material for embedded bioprinting^67^, a process when a print nozzle translates through a support material to deposit biomaterial inks^68^. A biomaterial ink formed from a spherical-particle based hydrogel was formed by centrifuging small (Figure 5A), gelatin microgels that melted upon heating to 37 °C. This ink was printed by extrusion into a fiber-based hydrogel that was contained withing a center well on a PDMS device (Figure 5B). To demonstrate the potential of the fiber-based hydrogel to support the printing of complex structures, a trifurcating and rejoining structure was printed (Figure 5B) with filament widths of ∼400 µm. Next, to show the fiber-based hydrogel could support a perfusable channel, we printed a single filament with a gelatin microgel ink containing a green fluorescent dye (Figure 5C) within the center well of the PDMS device. Two additional wells (not shown) are located on each side of the device to allow fluid to be perfused into the center well through printed channels when they connect to those wells. Whetn the gelatin ink was then melted by heating to 37 °C to leave a hollow channel, we could perfuse medium containing fluorescent microspheres into the channel. Interestingly, these channels remained open and perfusable without the need for interparticle crosslinking in experiments using PBS as the perfusing medium. Fiber-based hydrogels thus are support printing methods that enable the definition of complex tissue-like features within dynamic 3D environments, paving the way for future studies building microphysiological systems that can support cellular activities, including self-organization or growth or engineering vascularized tissue constructs.

**Figure 5.**
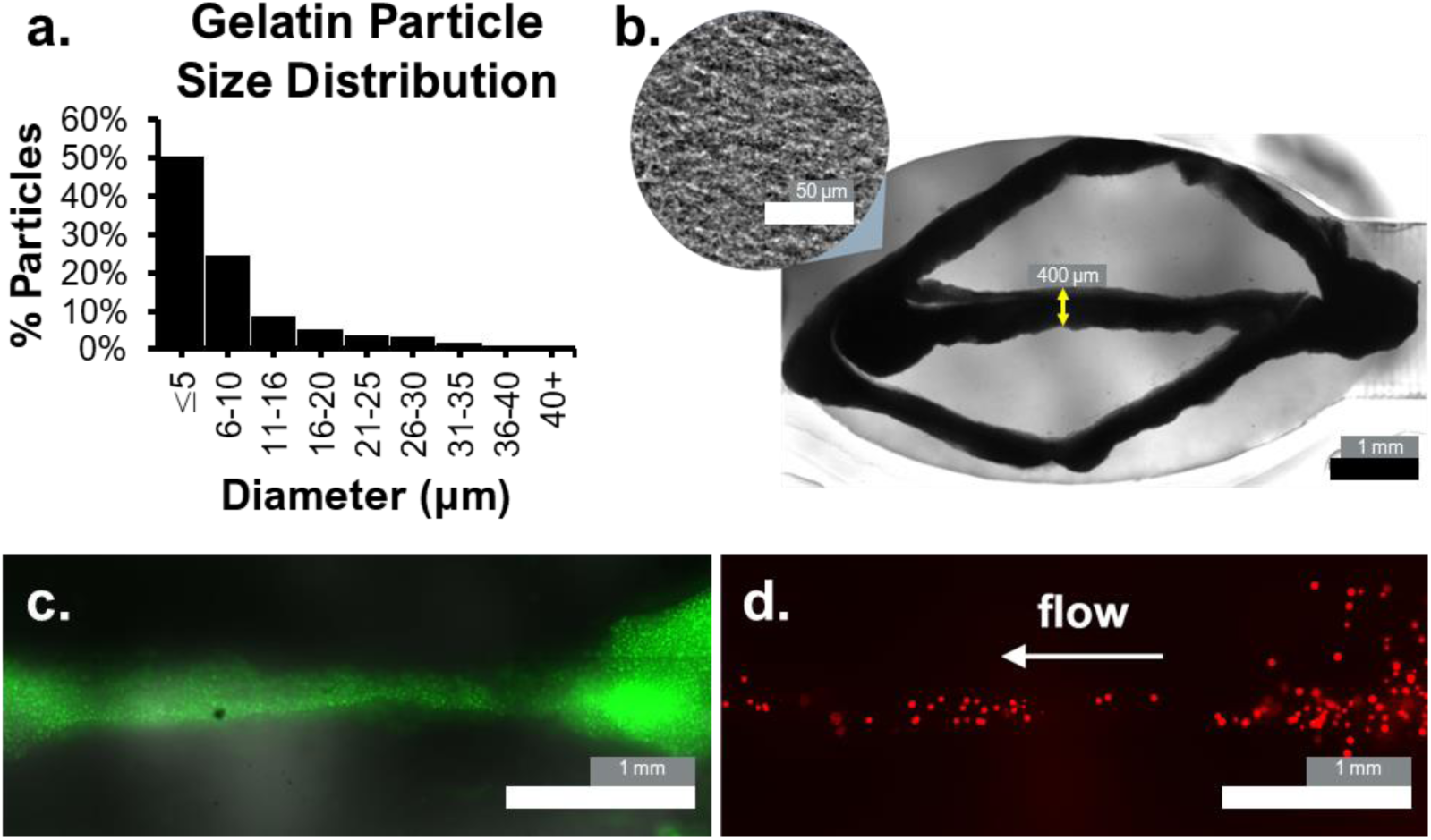
Embedded printing into fiber-based hydrogel supports using a gelatin microparticle biomaterial ink. (a) Size distribution of gelatin microparticles (spherical) that compose the biomaterial ink. (b) Embedded printing of trifurcating and rejoining filaments formed from gelatin microparticle infiber support (gray). Inset shows magnified image of microfiber support. Filament diameter (yellow arrow) is 400 µm. Scalebars: 1 mm and 50 µm (inset). (c) A printed filament of gelatin microgels containing green fluorescent marker prior to removal by melting. (d) A perfusable channel is formed after the gelatin microgels are melted. Here, the channel is being perfused with fluorescent (red) microspheres.

## IV. Conclusions and Future Outlook

Our work shows that fiber-based granular hydrogels created from high aspect ratio (∼15) electrospun PEG-based microfibers present unique properties as platforms for designing dynamic biomaterial systems. Viscoelasticity and stress relaxation are important characteristics of natural tissue but difficult to engineer into traditional 3D bulk hydrogel scaffolds. Particle-based systems offer the potential to dynamically respond to applied stress and strain to ultimately support dynamic cellular behaviors, but here we observed that particle-based hydrogels formed from microfibers exhibit tunable, ECM-mimetic properties that more closely mimic native tissue than particle-based hydrogels formed from spherical micorgels. The increased length:diameter ratio in fiber-based hydrogels enables long-range interactions between discrete fibers that enable higher yield strains for fiber-based granular hydrogels when compared to spherical-based systems with matched volume and matched dimension. Additionally, the fibrous composition gives rise to distinct scaffold properties, include fast and continuous stress relaxation in response to applied strains. Fiber-based hydrogels display a stress relaxing behaviors within cell-relevant strain regimes. With T_1/2_ values in the range of 1-100 s, stress relaxation occurs on timescales that are physiologically relevant for many tissue types, thereby offering user-defined design control over the time-dependent mechanics of the tissue culture scaffold.

Looking forward, the subcellular length scale diameters of fibers within the bulk of a fiber-based hydrogel might offer a more permissive 3D environment compared to spherical particles that are commonly sized to be on the same order of magnitude as cells, or larger. These materials’ dynamic properties also support embedded printing, presenting opportunities to design complex, heterogeneous architectures in biomaterials that can be designed with unique, dynamic biophysical properties. While the focus of this study was to characterize fiber-based hydrogels’ range of physical properties and compatibility with embedded printing, we expect that this class of granular hydrogels will facilitate work where dynamic behaviors are designed in defined systems for *in vitro* and *in vivo* applications.

