## Supporting information for "Fiber-based hydrogels for designing viscoelastic responses in particle-based biomaterials that support embedded 3D printing"

**Table of Contents**

**Figure S1. MALDI-TOF spectrum of GCDDD-FAM peptide**
**Figure S2. PEGVS fiber quantification**
**Figure S3. Effect of continuous phase concentration (dextran) on resultant microgel diameter**
**Figure S4. Effect of spin rate on resultant microgel diameter**
**Figure S5. Effect of disperse phase concentration (PEGNB and PEGSH) on resultant microgel diameter**

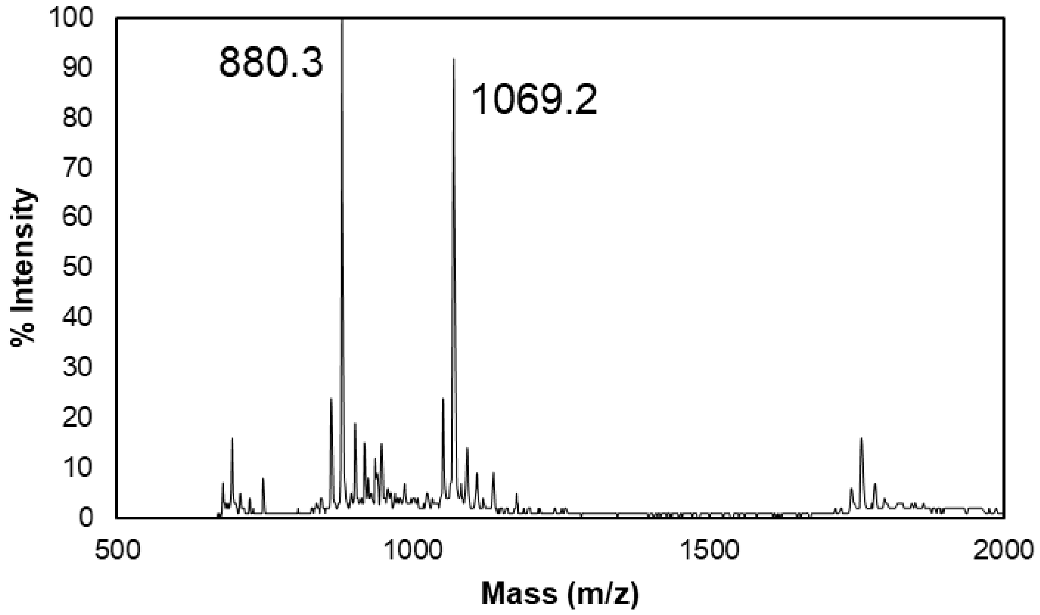


**Figure S1. MALDI-TOF spectrum of GCDDD-FAM peptide.** Confirmation of fluorescent peptide synthesis. Expected molecular weight: 881.8 Da; MALDI-TOF molecular weight: 880.3 Da and 1069.2 Da. 880.3 peak suggests successful synthesis, with some undesired side products (most notably, 1069.2 peak). We determined that since the 880.3 peak was the highest in magnitude, that synthesis was sufficient for use in this study.

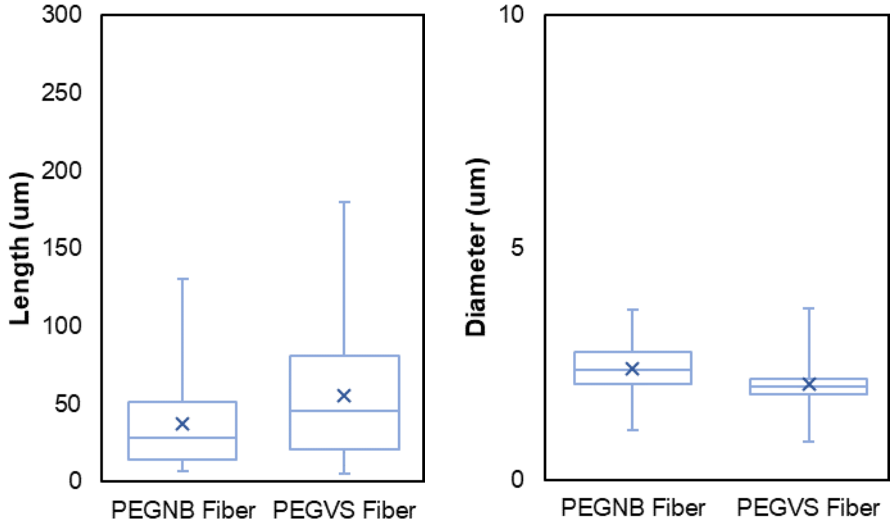


**Figure S2. PEGVS fiber quantification.** Comparisons between PEGNB fibers and PEGVS fibers show marginal differences in length and diameter. However, we determined that these values were sufficiently close to be considered interchangeable for this study – especially at the low concentrations (% v/v) of PEGVS fibers in the PEGNB fiber-based granular hydrogels (0-10% v/v).

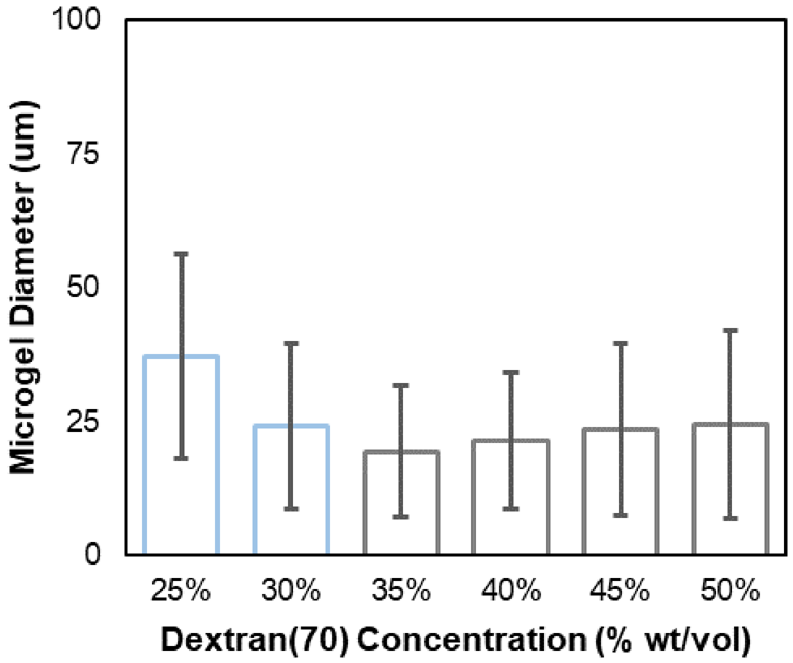


**Figure S3. Effect of continuous phase concentration (dextran) on resultant microgel diameter.** To determine the effect of the continuous phase concentration, the following variables were held constant: 6% w/v PEGNB, 4.2% w/v PEGSH (corresponding to [-SH]:[-NB]=0.7), and 800 RPM stir rate. We originally observed decreasing microgel diameter as dextran(70) concentration was increased from 25% to 30% w/v, then we saw a marginal increasing trend in diameter with increasing dextran(70) concentration above 30% w/v. This is likely due to the disperse phase aggregating in lower viscosity dextran(70) solutions to form larger droplets, with increasing viscosity (i.e., higher dextran(70) concentrations) supporting larger independent disperse phase droplets. Importantly, 25% w/v was the chosen continuous phase concentration for spheres (D) used in this study and 40% w/v was utilized for spheres (V).

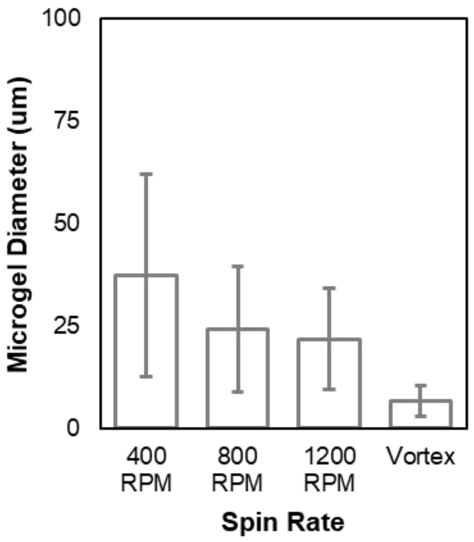


**Figure S4. Effect of spin rate on resultant microgel diameter.** To determine the effect of the spin rate, the following variables were held constant: 6% w/v PEGNB, 4.2% w/v PEGSH (corresponding to [-SH]:[-NB]=0.7), and 30% dextran(70). Overall, microgel diameter decreased with increasing spin rate. 800 RPM was utilized for spheres (D) in this study and spheres (V) were fabricated via vortexing at maximum speed rather than using a stir plate.

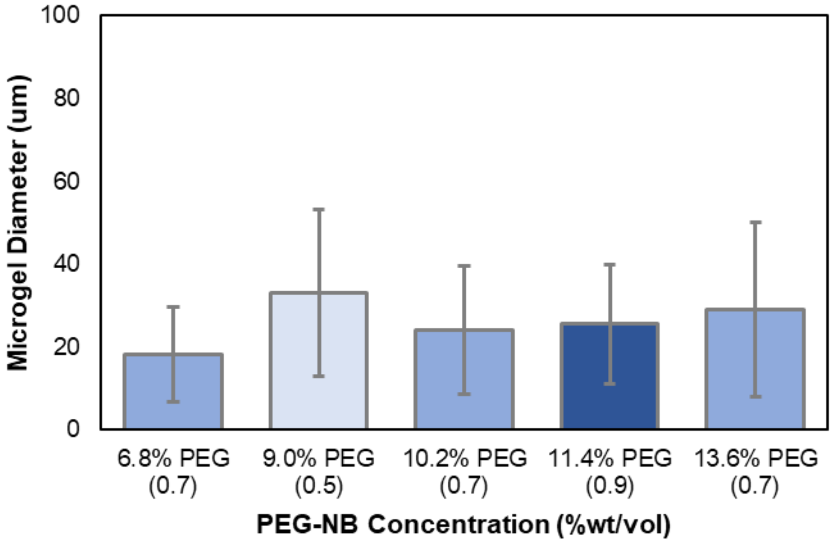


**Figure S5. Effect of disperse phase concentration (PEGNB and PEGSH) on resultant microgel diameter.** To determine the effect of the disperse phase concentration, the following variables were held constant: 30% dextran(70) and 800 RPM stir rate. The concentrations shown in the figure correspond to total % w/v between PEGNB and PEGSH, with the [-SH]:[-NB] ratio indicated below. Generally, increasing PEG concentration resulted in marginally larger microgel diameters, except for 9% PEG with [-SH]:[-NB]=0.5, which had the largest diameter due to the lowest crosslinking density allowing for the greatest swelling. 10.2% PEG – 6% PEGNB and 4.2% PEGSH – was utilized in this study for all microparticles.
